# Eye movements elevate crowding in idiopathic infantile nystagmus syndrome

**DOI:** 10.1101/2021.01.16.426927

**Authors:** Vijay K Tailor, Maria Theodorou, Annegret H Dahlmann-Noor, Tessa M Dekker, John A Greenwood

## Abstract

Idiopathic infantile nystagmus syndrome is a disorder characterised by involuntary eye movements, which leads to decreased acuity and visual function. One such function is visual crowding; a process whereby objects that are easily recognised in isolation become impaired by nearby flankers. Crowding typically occurs in the peripheral visual field, though elevations in foveal vision have been reported in congenital nystagmus, similar to those found with amblyopia. Here we examine whether the elevated foveal crowding with nystagmus is driven by similar mechanisms to those documented in amblyopia - long-term neural changes associated with a sensory deficit - or by the momentary displacement of the stimulus through nystagmus eye movements. We used a Landolt-C orientation identification task to measure threshold gap sizes with and without either horizontally or vertically placed pairs of flanking Landolt-Cs. Because nystagmus is predominantly horizontal, crowding should be stronger with horizontal flankers if eye movements cause the interference, whereas a sensory deficit would more likely be equivalent for the two dimensions. Consistent with an origin in eye movements, we observe elevations in nystagmic crowding that are above that of typical vision, and stronger with horizontal than vertical flankers and not found in amblyopic or typical vision. We further demonstrate the same pattern of performance can be obtained in typical vision with stimulus movement that simulates nystagmus. Consequently, we propose that the origin of nystagmic crowding lies in the eye movements, either through image smear of the target and flanker elements or through relocation of the stimulus into peripheral retina.

## Introduction

Our eyes are in constant motion. For some people, this motion is exaggerated and uncontrollable, a condition known as nystagmus (Papageorgiou, McLean, & Gottlob, 2014). Infantile Nystagmus Syndrome (INS) is a congenital condition with a typical onset prior to 6-months of age, which can be idiopathic (i.e. of no known cause) or related to a visual afferent abnormality - retinal dystrophies, albinism, low-vision, visual deprivation or a plethora of neurological conditions (Papageorgiou, McLean, & Gottlob, 2014). INS has an incidence of 14 per 10,000 population, with Idiopathic Infantile Nystagmus Syndrome (IINS) estimated at 1.9 per 10,000 (Sarvananthan et al., 2009). Nystagmus eye movements are most pronounced in the horizontal relative to the vertical plane (Abadi & Bjerre, 2002), with small torsional oscillations (Averbuch-Heller et al., 2002). Areas of visual function that are often reduced with nystagmus include visual acuity (Abadi & Bjerre, 2002), stereo-acuity (Guo, Reinecke, Fendick, & Calhoun, 1989; Ukwade & Bedell, 1999) and contrast sensitivity (Dickinson & Abadi, 1985). Particularly disruptive for foveal vision are the elevations in crowding, whereby objects that are easily recognised in isolation become impaired by nearby flankers (Chung & Bedell, 1995; Pascal & Abadi, 1995).

Crowding is a phenomenon that occurs in the typical peripheral visual field, disrupting the identification but not the detection of a target stimulus in clutter (Levi, Hariharan, & Klein, 2002b, 2002c; Pelli, Palomares, & Majaj, 2004). These disruptions occur over and above acuity limitations – an object can be large enough to see in isolation and yet still be difficult to recognise once flanked (Pelli, Palomares, & Majaj, 2004). The spatial extent of crowding can be quantified by measuring the transition point (or critical spacing) between largely correct and incorrect target identification. Bouma (1970) found this spacing to be 0.5× the target eccentricity – for example, a target at 6° eccentricity would be crowded by objects up to 3° away. This gives large spatial extents for crowding in peripheral vision (Toet & Levi, 1992), particularly in comparison to foveal crowding, where estimates of critical spacing range from 1-5 minutes of arc, depending on methodology (Flom, Heath, & Takahashi, 1963; Liu & Arditi, 2000; Coates, Levi, Touch, & Sabesan, 2018).

In peripheral vision, the spatial extent of crowding shows a number of variations in both size and shape. For instance, flankers positioned outwards from the target (with respect to fixation) have a greater effect on identification than inward flankers (Bouma, 1970). Toet and Levi (1992) further demonstrated a radial-tangential anisotropy, where the critical spacing is greater for flankers along the radial dimension compared to the tangential dimension relative to fixation. Variations have also been observed around the visual field, including the upper-lower anisotropy where crowding is stronger in the upper compared to the lower visual field (Petrov & Meleshkevich, 2011; Greenwood, Szinte, Sayim, & Cavanagh, 2017). In contrast to peripheral vision, the spatial extent of foveal crowding has been found to be more circular, with equivalent crowding in the horizontal and vertical dimensions. This has been observed in typical vision using various stimulus configurations (Flom, Heath, & Takahashi, 1963; Toet & Levi, 1992; Pluháček, Musilová, Bedell, & Siderov, 2021) and likewise in the amblyopic fovea (Levi & Carney, 2011).

Elevations in foveal crowding occur in both children and adults with strabismic amblyopia (Levi & Klein, 1985; Levi, 2008; Greenwood et al., 2012). This developmental disorder occurs through a misalignment in the visual axis, which can lead to a reduction in acuity with the deviating eye (McKee, Levi, & Movshon, 2003; Barrett, Bradley, & McGraw, 2004). Amblyopic crowding exhibits many of the same attributes as peripheral crowding (Levi, Hariharan, & Klein, 2002c), with commonalities in the nature of crowded errors suggestive of a common mechanism. In peripheral vision, crowding produces systematic errors that follow either an average of the target and flanker features (Parkes et al., 2001; Greenwood, Bex, & Dakin, 2009) or substitution of the flankers (Ester, Zilber, & Serences, 2015). A population-coding model that inappropriately combines target and flanker responses can account for both error types (Harrison & Bex, 2015). Averaging and substitution errors have similarly been found in the amblyopic fovea, with a population-coding model again able to reproduce these errors (Flom, Weymouth, & Kahneman, 1963; Kalpadakis-Smith, Tailor, Dahlmann-Noor, & Greenwood, 2017). Amblyopic elevations have been linked to a sensory deficit with a reduction in the number of neurons driven by the amblyopic eye (Kiorpes & McKee, 1999) and increases in population receptive field size (Clavagnier, Dumoulin, & Hess, 2015) observed in cortical areas V1 and above.

Although the elevation of foveal crowding in nystagmus has been clearly demonstrated (Chung & Bedell, 1995; Pascal & Abadi, 1995), the origin of this deficit is less clear. Pascal and Abadi (1995) measured acuity using oriented Landolt-Cs, with crowding induced using flanker bars. Crowding was elevated in individuals with idiopathic nystagmus and albinism relative to controls, though the difference was only significant for the idiopaths. The authors attributed the difference in these elevations to variations in the pattern and amplitude of eye movements between idiopathic and albinism groups, with the implication that image motion caused by eye movements may be the primary determinant of nystagmic crowding. Chung and Bedell (1995) similarly examined acuity with Landolt-C targets and induced crowding with a black or white surround, finding elevations in both the magnitude and spatial extent of crowding in several observers with nystagmus compared to controls. To test the role of stimulus motion on these crowding effects, Chung and Bedell (1995) applied a simulated nystagmus movement to the stimulus, which reduced acuity and elevated crowding in controls. However, these simulated deficits did not reach the same level of impairment as the nystagmus participants, suggesting that eye movements alone are insufficient and that an underlying sensory deficit causes the remainder of the deficit. Together, these findings present evidence both for a long-term sensory deficit and for more momentary effects of eye movements, such as image smear.

Studies of other performance decrements in nystagmus reveal a similarly mixed picture. Most widely studied in this context is the decrease in visual acuity associated with nystagmus (Cesarelli, Bifulco, Loffredo, & Bracale, 2000), which has been shown to correlate with the amount of time spent with the stimulus under *foveation* (Abadi & Worfolk, 1989; Dell’Osso, van der Steen, Steinman, & Collewijn, 1992; Bedell, 2000; Dell’Osso, 2002; Dell’Osso & Jacobs, 2002; Theodorou, 2006), a property that is met when the fovea is close to the stimulus during a period of slow eye movements. An effect of eye movements is also apparent in the anisotropic pattern of elevation that has been observed in orientation discrimination tasks (Abadi & King-Smith, 1979; Ukwade, Bedell, & White, 2002; Dunn et al., 2014) where performance is better for horizontally-than vertically-oriented lines. Because vertically oriented lines would smear into each other to a greater extent with the predominantly horizontal eye movements, this anisotropy is consistent with an origin in eye movements. However, the observation of these deficits under tachistoscopic conditions, which would minimise image smear, suggests that a longer-term sensory deficit may arise (Abadi & King-Smith, 1979; Ukwade, Bedell, & White, 2002; Dunn et al., 2014). Similar anisotropies are also found for bisection acuity (Ukwade & Bedell, 2012), with greater elevation for horizontal judgements than vertical in individuals with congenital nystagmus (again consistent with the predominant direction of image smear). However, thresholds for *both* horizontal and vertical elements were elevated relative to control participants, suggesting that at least part of the impairment in visual function may derive from a sensory deficit, as in amblyopia.

Given the divergent conclusions of these studies, we do not currently know whether the elevated crowding in nystagmus originates from the momentary image smear caused by eye movements, an underlying sensory deficit, or both. Here, we sought to investigate these underlying mechanisms. To do so, we measured the spatial extent of crowding in idiopathic infantile nystagmus with flankers placed either horizontally or vertically relative to a target Landolt-C element. Because nystagmus eye movements are predominantly horizontal, factors related to these eye movements (e.g. image smear) should cause horizontally-placed flankers to be more disruptive than vertical flankers. If nystagmic crowding derives from momentary eye movements, we should therefore find a horizontal elongation of the spatial extent of crowding. In contrast, if nystagmic crowding derives from a sensory deficit associated with factors such as an increase in receptive field size, then the elevation in crowding should be equivalent for the two dimensions, as observed in amblyopia (Levi & Carney, 2011). A third possibility is that both momentary interference and longer-term deficits are present, in which case thresholds should be elevated with both horizontal and vertical flankers but to a larger extent with the horizontal flankers.

In order to examine the effect of nystagmus in participants with no observable retinal or neural defects, we examined those with idiopathic infantile nystagmus. In Experiment 1, we compared this group to adults with strabismic amblyopia, a population where crowding is more clearly derived from a sensory deficit, as well as participants with typical vision. We then examine the origins of nystagmic crowding by applying nystagmus image motion to crowded stimuli viewed by participants with typical vision in Experiment 2.

## Experiment 1: Crowding with horizontal and vertical flankers

### Methods

#### Participants

Thirty-one adults underwent a full orthoptic examination to ensure they met inclusion and exclusion criteria into one of three clinical groups: nystagmus (n = 8, M_age_ = 30.3 years), strabismic amblyopia (n = 10, M_age_ = 36.2 years) or controls (n = 10, M_age_ = 32.1 years). All participants were between the ages of 19-49 years old with no neurological conditions. Control participants had to achieve a best corrected visual acuity (BCVA) in each eye of 0.20 logMAR or better, with no strabismus or nystagmus present. In the amblyopic group, participants needed to demonstrate strabismic or combined strabismic and anisometropic amblyopia with manifest strabismus, as well as a BCVA difference of 0.20 logMAR between the two eyes, a BCVA in the amblyopic eye between 0.20 and 1.00 logMAR, and a BCVA of 0.20 logMAR or better in the fellow eye. For the nystagmus group a horizontal, vertical or torsional nystagmus waveform had to be present with a diagnosis of idiopathic nystagmus (without visual afferent abnormality), a BCVA of 1.00 logMAR or better and no manifest strabismus. Three adults with nystagmus were excluded due to an incorrect nystagmus diagnosis and are not included in the above tally.

Clinical characteristics can be found in Appendix A. BCVA was measured using a logMAR chart, with a mean visual acuity (±1 SD) for the control group of - 0.09 ±0.11 logMAR, 0.56 ±0.28 logMAR for the amblyopic group, and 0.29 ±0.26 logMAR for the nystagmus group. Controls all demonstrated excellent stereo-acuity (measured with the Frisby stereo-test), with no ocular motility imbalances. Amblyopes all had a predominantly horizontal strabismus, with 2/10 exhibiting a small vertical component. Nine of the amblyopes demonstrated no stereopsis, with one exhibiting stereo-acuity at 170 seconds of arc. Nystagmats all showed nystagmus eye movements that were predominantly horizontal in direction, and which involved either a jerk or pendular movement. No strabismus was present and stereopsis was found in all cases, with a mean stereo-acuity of 350 seconds of arc.

#### Apparatus

Testing was undertaken at Moorfields Eye Hospital, London. Experiments were programmed using Matlab (The Mathworks, Ltd.) on a Dell PC running PsychToolBox (Brainard, 1997; Pelli, 1997). Stimuli were presented on an Eizo Flexscan EV2736W LCD monitor, with 2560 × 1440-pixel resolution, 60Hz refresh rate, and a physical panel size of 59.7 × 33.6 cm. The monitor was calibrated using a Minolta photometer, with luminance linearized in software to give a maximum of 150 cd/m^2^. Participant responses to stimuli were indicated by a keypad. An EyeLink 1000 (SR research, Ottawa, Canada) recorded the position of the dominant eye at a sampling rate of 1000 Hz. Participants had their head positioned on a chin rest with a forehead bar to minimise head movement. Stimulus presentation was binocular for nystagmus participants and monocular for the others, achieved through occlusion of either the non-amblyopic eye for amblyopes or the non-dominant eye for controls. Viewing distance was varied between participants based on their BCVA in the orthoptic examination: 7 were tested at 78cms (2 nystagmats, 5 amblyopes), 7 at 150cms (4 nystagmats, 3 amblyopes), 1 at 200cms (a nystagmat) and 13 at 300cms (1 nystagmat, 2 amblyopes, 10 controls). This ensured both adequate resolution to measure acuity and sufficient range on screen to measure the extent of crowding. All participants wore their refractive correction where required, with no correction made for presbyopia. Data were analysed in Matlab and SPSS.

#### Stimuli and procedures

Target and flanker stimuli were Landolt-C letters presented at the centre of the screen, either in isolation (unflanked) or flanked by two Landolt-C elements positioned either horizontally or vertically. Elements were presented at 99.6% Weber contrast against a mid-grey background (Figure 1). Participants identified the position of the gap of the Landolt-C (four alternative forced choice, 4AFC). The orientation of the Landolt-C was always oblique, either at 45°, 135°, 225°, or 315°. These oblique orientations were selected following pilot testing (by authors VKT and JAG) with gap orientations at cardinal positions (0°, 90°, 180° and 270°), where we found improved identification of orientation when target gap orientations were orthogonal to the flankers (e.g. up/down gap orientations were better recognised when horizontal flankers were present and vice versa). On modification of the orientation to oblique positions we found the identification of gap orientation to be equivalent across all configurations. Given evidence that nystagmats are “slow-to-see” (Hertle et al., 2002; Wang & Dell’Osso, 2007; though cf. Dunn, Margrain, Woodhouse, & Erichsen, 2015), stimuli were presented with unlimited duration until a response was made, at which point they were removed from the screen. This ensured that any performance decrements in crowded conditions were not simply due to suboptimal presentation times. Fixation guides were present throughout the experiment in the form of 4 white lines (the opposite polarity to the target) at a Weber contrast of 74.7%, located at the cardinal positions with a length of 50 pixels and with the inner edge separated from fixation by 15 times the stimulus diameter to avoid overlap. A 500ms inter-trial interval with a blank screen (leaving only a fixation point and the fixation guides) was then presented prior to the next stimulus. Feedback was not given. Participants were encouraged to make a choice in a timely manner.

**Figure 1.**
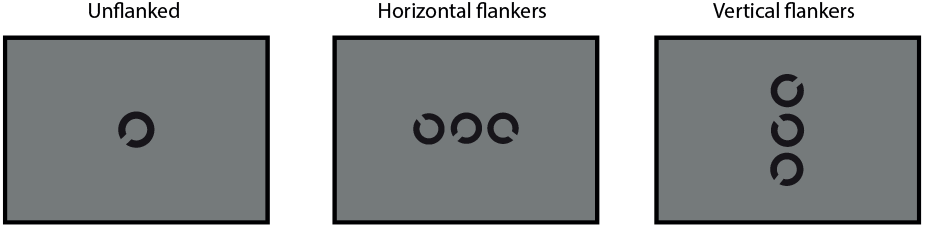
Examples of the Landolt-C discrimination task, unflanked (single presentation of a Landolt-C), flanked-horizontal (participants choose the orientation of the middle Landolt-C, flanked horizontally by two other Landolt-C elements) and flanked-vertical.

The gap size and stroke width of target and flanker Landolt-C elements was 1/5^th^ of the diameter, which was scaled using an adaptive QUEST procedure (Watson & Pelli, 1983) that converged on 62.5% correct (midway between chance and 100% correct). To avoid rapid convergence of the QUEST we added variance to the gap sizes presented on each trial by adding a value selected from a Gaussian distribution (with 0 mean and 1 SD) multiplied by 0.25 of the current trial estimate of the threshold. This minimised the number of trials presented at the same size in order to improve the subsequent fit of psychometric functions to the data (Kalpadakis-Smith et al., 2018). When flankers were present, their size matched the target, with a centre-to-centre separation from the target of 1.1 times their diameter, following Song, Levi, and Pelli (2014) who found this to be the ideal spacing to measure crowding effects in normal peripheral vision and the amblyopic fovea. With this scaling we can see the effect of crowding as an elevation in threshold gap size in flanked conditions relative to performance with an unflanked target. The constant scaling of both stimulus size and flanker spacing means we can then calculate the spatial extent of crowding as gap size × 5 (stimulus diameter) × 1.1 (spacing).

Participants commenced with 5 practice trials (identifying the orientation of the gap of the target Landolt-C) at the start of each block of trials, which were not included in the main analysis. Each block consisted of 65 trials, including the 5 practice trials, with 4 repeats per block to give a total of 240 trials (excluding practice) for each stimulus condition (unflanked, horizontal flankers and vertical flankers). The whole experiment took approximately 2 hours, split over 1-2 sessions.

All procedures were approved by the NHS North Thames Research Ethics Committee. Participants were reimbursed for their time and travel expenses.

#### Eye tracking

Eye movements were recorded with the EyeLink 1000 to characterise the nystagmus eye movements and to monitor participant gaze throughout the trials. Calibration was undertaken at the beginning of each block of trials, with amblyopic and control participants calibrated using the EyeLink 1000 inbuilt 5-point calibration routine. Calibration for amblyopes was performed with the amblyopic eye and the dominant eye in the controls.

Several of the participants with nystagmus could not undertake the inbuilt EyeLink calibration process due to the large variability in their eye position. The nystagmus participants were thus calibrated using a novel 5-point system, similar to several procedures reported recently for the calibration of participants with nystagmus (Dunn et al., 2019; Rosengren, Nyström, Hammar, & Stridh, 2020). Prior to this calibration, an observer with typical vision undertook the inbuilt EyeLink calibration, which allowed an approximation of eye position for the participants with nystagmus. For our custom calibration, white circles were then presented as fixation targets with a diameter of 0.5° and a brightness of 150 cd/m^2^. These were presented binocularly in a random order at the centre of the screen and separated by 5° at the following positions – 0°, 90°, 180°, 270°. The participant was instructed to look at the fixation target for 5 seconds and move to the new location. EyeLink recordings were observed by the examiner and if a loss of recording or excessive blinking was detected the calibration was repeated.

Calibration recordings were used to perform a post-hoc calibration of eye position using a geometric transformation. Data from the first 0.5 seconds of each calibration trial location was discarded to allow for fixation to arrive on the target. Time points where the velocity of the eye was more than ±1SD from the mean velocity of the eye across the whole trial were removed, leaving only the positions of the eye where the velocity was slow. The mean of these slow phases became our fixation locations for each of the five calibration target locations. These values were compared to the calibration target locations on the screen, with an affine geometric transformation used to align the two sets of points. This transformation was chosen after pilot testing as best able to preserve the ratios of the distances between the points lying on a straight line to give the final geometric transformation value. This transformation was then applied to the eye positions recorded within the main trials of the experiment. Each block of trials had a corresponding calibration file as head position may have varied between blocks. Difficulties were found with one nystagmus participant where one calibration file was unable to constrain the affine geometric transformation value. To overcome this, we applied the affine geometric transformation from a prior block of trials.

Once the affine geometric transformation was calculated and applied to all corresponding trials the final processing of the eye fixation was undertaken. Blinks were removed following identification through absent pupil readings. Blink generation was also removed by identifying readings within 50 milliseconds before and after the blink, as determined during pilot testing. For all participants, eye positions within each trial were converted to error values around the fixation in degrees of visual angle. At each timepoint, velocity was computed as a moving estimate of 3 successive eye-position samples in order to reduce noise within the velocity calculations (Engbert & Kliegl, 2003).

### Results

#### Behavioural results

Blocks of trials were combined for each participant and stimulus condition (unflanked, horizontal flankers and vertical flankers), to give 240 trials per stimulus condition. For each stimulus size that was presented, the corresponding proportion correct scores for responses were then collated. Psychometric functions were fitted to the behavioural data for each stimulus condition using a cumulative Gaussian function with 3 free parameters (midpoint, slope and lapse rate). Because the variability added to the QUEST gave variable trial numbers for each gap size, this fitting was performed by weighting the least-squared error value by the number of trials per point. Figure 2A plots example proportion correct values for one nystagmus participant in the unflanked condition along with the best-fitting psychometric function (black solid line). Figure 2B and 2C show data for the horizontal and vertical flanker conditions. Note that with the variable number of trials for each gap size that some proportion correct values can lie at the extremes due to a low number of trials (e.g. the lowest value in Figure 2B, which derives from a single trial), but that the weighted function fitting de-emphasises these values. Gap-size thresholds were derived from the psychometric function when performance reached 62.5% correct (mid-way between chance and ceiling) represented by the black dotted line, and converted to degrees of visual angle.

**Figure 2.**
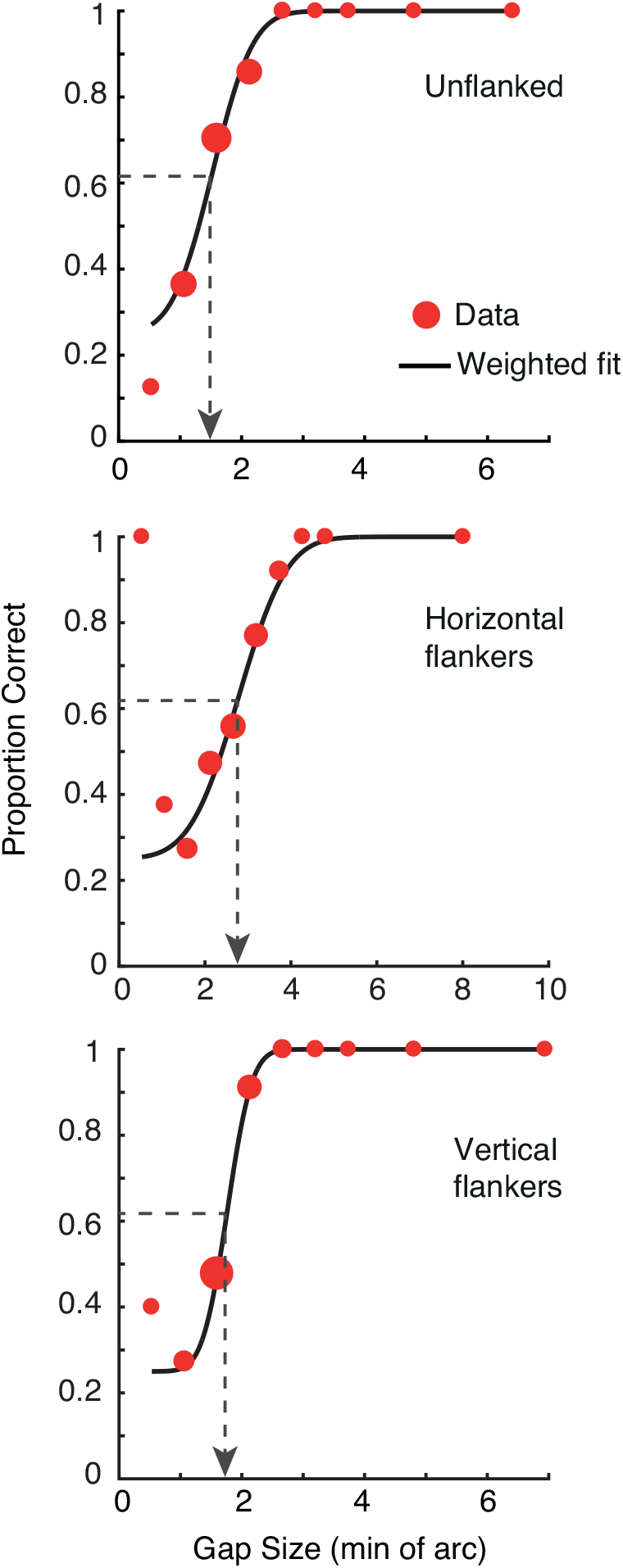
A-C. Example data for the 3 stimulus conditions (unflanked, horizontal flankers and vertical flankers) in a patient with nystagmus. Red circles plot the proportion of correct responses at each of the gap sizes presented, with dot size indicating the relative number of trials at each gap size. The black solid line plots the best-fitting psychometric function. Thresholds were taken at 62.5% correct, shown as the black dashed line and its corresponding threshold on the x-axis. Note the different scales along the x-axes.

Figure 3A plots the mean gap thresholds in minutes of arc (left y-axis) for all participant groups, along with logMAR equivalent values (right y-axis). In the control group, gap thresholds were low overall in all stimulus conditions with the unflanked condition (green bar) having the lowest threshold and rising slightly with either horizontal flankers (blue bar) or vertical flankers (red bar). The amblyopic group demonstrated large elevations of gap thresholds in all tasks compared to the control group, particularly when crowded. Finally, thresholds for the nystagmus group were elevated in all conditions compared to the control group, though to a lesser extent than the amblyopic group

**Figure 3.**
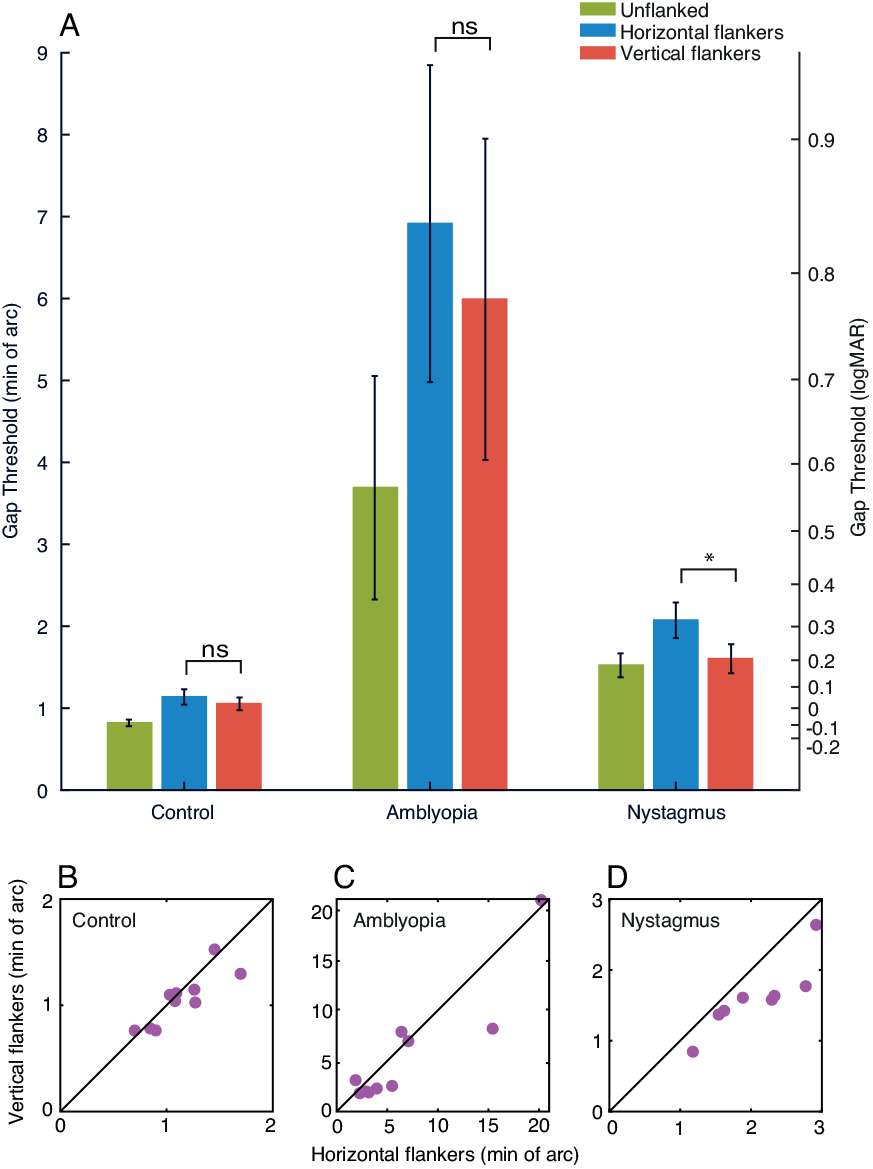
A. Gap thresholds for all participant groups, plotted in minutes of arc (left y-axis), with comparison to their logMAR equivalents (right y-axis). The blue bars plot thresholds for the unflanked condition, green bars the horizontal flanker condition and red bars the vertical flanker condition. Error bars represent the SEM, with *= significant and ns = not significant. B-D. Gap thresholds (in min. of arc) with horizontal flankers for each individual plotted on the x-axis, against gap thresholds with vertical flankers on the y-axis. The black line represents perfect correspondence between gap thresholds with horizontal and vertical flankers. Purple circles represent individual participants. Note the variation in X and Y scales.

We undertook a 3×3 mixed effects ANOVA to further examine these differences, with factors for participant group and stimulus condition. LogMAR values were used here (calculated for each individual threshold) to reduce heteroscedasticity in the data. A main effect of stimulus condition was found (F_(2,50)_ = 37.700, p<0.001), indicating that the presence of flankers affected the gap thresholds. The main effect of participant group was significant (F_(2,25)_ = 18.330, p<0.001, and there was a significant interaction between stimulus condition and participant group (F_(4,50)_ = 5.031, p=0.002). This indicates that the effect of stimulus condition on gap thresholds differed for the participant groups, which we examined with a series of a priori contrasts.

To test between the image motion or sensory deficit hypothesis regarding nystagmic crowding, we undertook paired sample t-tests comparing horizontal and vertical flanker conditions for each group, again in logMAR units. As shown in Figure 3A, there is no clear difference between horizontal flanker and vertical flanker thresholds in the control group and indeed this difference was not significant, t_(9)_=1.823, p=0.102. Although the amblyopic group show higher horizontal flanker thresholds compared to vertical flanker thresholds on average, this difference was not significant, t_(9)_=1.610, p=0.142. In contrast, for the nystagmats there is a clear difference in performance, with the horizontal flankers producing higher thresholds than vertical flankers, which was found to be significant t_(7)_=5.277, p=0.001.

To further explore these differences, Figure 3B-D plots the relationship between the horizontal flanker and vertical flanker conditions for each individual. In this plot the black line represents a direct correspondence between the horizontal and vertical flanker thresholds. An individual with no difference between the two stimulus conditions would lie on this line, while an individual with an elevated horizontal flanker threshold would lie below the unity line, and vice versa. In the control group, all individuals were clustered around the unity line, with no consistent difference between the two crowding conditions – 5/10 observers demonstrated higher thresholds with horizontal flankers than with vertical. In the amblyopia group, 7/10 participants had higher thresholds in the horizontal flanker condition (below the line of unity) compared to the vertical flanker condition. In other words, although thresholds were worse on average with horizontal flankers than with vertical, this was not wholly consistent amongst the amblyopic participants. For the nystagmus group, all participants consistently had higher thresholds with horizontal flankers than in the vertical flanker condition, with all data points lying below the line of unity. This specific impairment for the horizontal flanker condition in the nystagmus individuals supports the findings in the paired sample t-test, both of which follow the prediction of image motion as the basis for nystagmic crowding. In other words, our findings reveal stronger crowding in the horizontal flanker condition than the vertical flanker condition in the nystagmus group, suggesting that nystagmic eye movements (which are predominantly horizontal in movement) could be the cause of elevations in horizontal crowding.

Because our scaling approach also allows estimation of the spatial extent of crowding (Song, Levi, & Pelli, 2014), we calculated these values in each of our three participant groups. In the control group, the extent of crowding was small and equivalent for both horizontal and vertical flankers, with mean values of 0.104° and 0.097° respectively. This increases markedly in the amblyopic group with mean values of 0.634° and 0.549°, though as above these values do not differ significantly. In the nystagmus group, spatial extent values were on average 0.190° and 0.147° with horizontal and vertical flankers, respectively.

#### The role of eye movements in crowding

The above results are consistent with eye movements being the cause for the elevated foveal crowding in nystagmus. Were this the case, the elevated thresholds in the horizontal flanker condition could occur through either image smear (where target and flankers’ smear into each other as the stimuli move across the retina) or the shift of stimuli into peripheral vision (relocation of the stimuli into peripheral retina where crowding is known to occur). To further consider this relationship, we examined eye movement properties for each group, and considered their potential influence on gap thresholds.

Analysis of the eye movement properties reveals that the eye position variability across all trials on the horizontal plane was lowest in the control group (mean of the standard deviation across trials = 0.56°), which increased for the amblyopes (0.99°), and further again for the nystagmus group (1.60°). On the vertical plane, eye position variability was similar to the horizontal variability for controls (0.67°) and amblyopes (0.95°). For nystagmats the variability was higher than controls and amblyopes (1.08°), but lower than the horizontal variability in their eye movements. The average velocity of the eye (including both steady fixation and microsaccades) across all trials was slowest in the control group (6.78°/sec), which increased in the amblyopic group (10.5°/sec), and further again in the nystagmus group (25.3°/sec). Note that these values are higher than some estimates of fixational stability and eye velocity (Collewijn, Martins, & Steinman, 1981; Chung, Kumar, Li, & Levi, 2015), likely due to the inclusion of microsaccades and slow drift in our measurements.

If nystagmic crowding is due to the image motion caused by these eye movements, then these properties of variability and velocity should correlate with gap-size thresholds. In other words, with increases in eye position variability and eye speed we should see an increase in the size of gap thresholds. Contrary to this, our analysis of the individual gap thresholds against the standard deviation of the position or variability in velocity found no such correlations (see Appendix B). These measures of position variability and velocity are however somewhat crude estimates of the relationship between eye movements and the stimuli shown on screen, given the variability of nystagmus eye movements in particular.

We next examined the effect of foveation duration in individual trials on performance. Foveation criteria were derived from prior studies (Dell’Osso & Jacobs, 2002), given the observation that better measures of acuity are associated with a higher frequency and longer duration of foveation periods (Abadi & Worfolk, 1989; Cesarelli et al., 2000). As above, foveation is defined as a period when fixation is within a restricted spatial window around the target *and* where the velocity of the eye is lower than a cut-off value. To apply this to our data, we used parameters from Dell’Osso (2002) who developed the eXpanded Nystagmus Acuity Function (NAFX), where foveation was determined using eye positions within ±0.5-4° of the target and with a velocity below 4-10°/sec. We evaluated different criteria within these ranges for position and velocity to define foveation and non-foveation periods, allowing us to establish the amount of time in each trial that met this range of foveation criteria. This allowed us to split trials into those with the longest foveation durations (‘best-foveation’), and those with the shortest (‘worst-foveation’), to determine whether foveation and gap thresholds are associated.

To explore these criteria, Figure 4A shows the average amount of time within each trial classed as foveation within the different spatial windows, as defined by the parameters of the NAFX (i.e. between ±0.5-4°). Here, velocity was set at the highest end of the NAFX criteria (10°/sec). Overall the control participants (red line) and amblyopes (green line) have comparable durations of time spent within the foveation window, both of which are longer in duration in comparison to the nystagmats (blue line), regardless of its size. Increasing the size of the spatial window gave a modest increase in foveation time, with all participant groups reaching a ceiling at a spatial window around 1.5-2°. Figure 4B shows the average duration of foveation within the different velocity windows, once again determined by the NAFX (4-10°/sec), with the spatial window set at 4°. Again, the average duration of foveation is greatest for the controls and amblyopes, and considerably lower for the nystagmats. Increasing the window and allowing faster eye movements to be included increases the duration of foveation for all groups, though the nystagmats always show the least foveation. Although the precise amount of foveation varies depending on the parameters that are applied, nystagmats consistently show the least foveation in a given trial.

**Figure 4.**
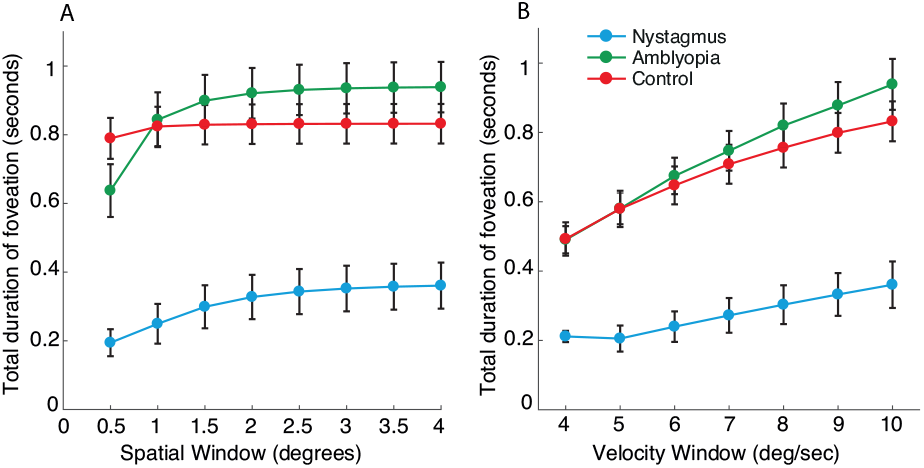
A. The average amount of time within each trial when the eye position falls within spatial windows of varying sizes around the target location (with a fixed velocity window), plotted separately for each of the participant groups. The X-axis shows the radius of the different spatial windows in 0.5° increments. The Y-axis is the total duration of time within the spatial window. B. The average duration of time spent where the velocity of the eye is within a range of foveation velocity windows (with a fixed spatial window). The X-axis shows the different velocity windows in 1°/sec increments, while the Y-axis shows the total duration of the trial within this window. Error bars represent the SEM.

We next divided these trials into best-foveation and worst-foveation categories, by first examining the distribution of foveation durations to find a combination of spatial and velocity windows that produced a normal distribution of durations across all trials, without any ceiling or floor effects. Methods and results can be found in Appendix C. If nystagmic crowding is caused by image motion and differences in foveation, we should find lower thresholds for best-foveation trials compared to the worst-foveation trials. However, when splitting the data in this way we find identical patterns of thresholds for all stimulus conditions, with significantly worse thresholds with horizontal compared to vertical flankers in both best- and worst-foveation conditions.

Overall, our results have shown that thresholds for the unflanked and flanked conditions are elevated in nystagmus compared to typical adults, albeit to a lesser extent than the elevations seen with amblyopia. These deficits could be due to a long-term sensory change or to the image motion induced by eye movements. The significant difference between the gap thresholds measured in horizontal and vertical flanker conditions in the nystagmus group but not in the amblyopes and controls is consistent with the hypothesis that these deficits are caused by image motion through the momentary eye position changes. However, the lack of a correlation between gap-size thresholds and either the variability in eye position or eye velocity, as well as the lack of effect of foveation on thresholds, means that we cannot rule out a sensory deficit to nystagmic crowding. Nonetheless, we note that the individual variations in these eye-movement properties (position, velocity, and foveation) are small relative to the differences at the group level, where we find large differences that could potentially account for the large differences in thresholds between groups. It remains possible that these large differences in eye-movement properties could cause the difference in crowding between the participant groups.

## Experiment 2: Simulation of nystagmic crowding in typical vision

If nystagmic crowding is caused by image motion then its horizontal elongation (where thresholds are worse with horizontal than vertical flankers) should be reproducible in typical adults when stimuli move on-screen in the same way as nystagmic eye movements. In Experiment 2 we measured thresholds in the same stimulus conditions as above for adults with typical vision, either with stationary stimuli or with stimulus motion derived from the eye-movement recordings of the nystagmus group from Experiment 1. If image motion is the basis for nystagmic crowding then we should see the same horizontal-elongation of crowding with this motion applied to the stimulus in adults with typical vision. On the other hand, if nystagmic crowding is derived from a sensory deficit then the simple application of stimulus motion should fail to reproduce the observed elevations in thresholds and the horizontal-elongation of crowding.

### Methods

#### Participants

Ten adults with typical vision (M_age_ = 31.4 years) were recruited, including one of the authors (JG). All participants wore full refractive correction if needed (M_BCVA_ = −0.10 logMAR) and were tested binocularly. No strabismus or nystagmus was present.

#### Stimuli and procedures

Stimulus conditions were as in Experiment 1 (unflanked, horizontal flankers and vertical flankers), with the same 4AFC Landolt-C orientation identification task. Presentation time was unlimited. Here, the 3 stimulus conditions were completed in 3 motion conditions. For the first ‘no motion’ condition, stimuli were static at the centre of the screen, as in Experiment 1. The latter two conditions had motion applied to the stimuli to simulate the pattern of nystagmus eye movements: in the motion-fixation condition the stimulus moved while participants fixated the centre of the screen, while for the motion-following condition participants were allowed to follow the stimulus as it moved. Fixation was maintained as described above in Experiment 1. Pilot testing revealed the motion-fixation condition to be more difficult than the motion-following condition; both were included here to allow measurement of performance at these different levels to see whether one might produce thresholds that better match nystagmic performance. Each block consisted of 60 trials and 5 practice trials as in Experiment 1, with each block repeated 3 times to give a total of 180 trials for each testing condition and motion condition. The same monitor, computer, and EyeLink setup from Experiment 1 were used. In the no motion and motion-fixation conditions, fixation was monitored more closely, with a tolerable fixation zone of 1.5 degrees radius around the centre of the screen. If fixation deviated from the central fixation zone, the trial was cancelled and repeated at the end of the block. In the motion-following condition, trials were not cancelled when the eye diverged from this zone, though a given trial would not commence until fixation was within the central fixation zone in order to stop anticipation of the stimulus motion.

#### Generation of the motion waveforms

In the two motion conditions, stimulus motion was derived from eye movements recorded from participants with nystagmus. These nystagmus waveforms were obtained from the 5-second central fixation recordings made during the 5-point calibration in Experiment 1. Because it has been shown that gaze position can alter the nystagmus waveform (Abadi & Whittle, 1991), only the central fixation recordings (and not the eccentric locations) were used.

Sixty waveforms were selected after visually inspecting their quality, with a preference for recordings where the waveform was repeated regularly and without any loss of recording. Both horizontal and vertical elements of the waveforms were used. The initial 500ms of all trials were removed to counter any re-fixation movements to the central fixation position, leaving us with 4.5 second recordings. Waveforms were looped to provide a continuous movement, with an additional 500ms section added to the end that joined the end of the waveform to the beginning to allow a continuous loop. This was done by linearly interpolating between the final eye position recording with the first recording. An example recording is shown in Figure 5A. Because the original waveforms were recorded at 1000Hz, they were down-sampled to match the screen refresh rate of 60Hz, by sampling the recordings at every 17^th^ interval (shown in Figure 5B). Once sampled in this way, the waveform was centred to pass through the central position of the screen by subtracting the mean position of the waveform. Centering the sampled waveform allowed us to ensure that the stimulus passed through the fovea at some point in the trial, along with the restriction of Y positions to within ±1° of the screen centre. Waveforms were smoothed by applying a three-timepoint boxcar average to the X and Y coordinates.

**Figure 5.**
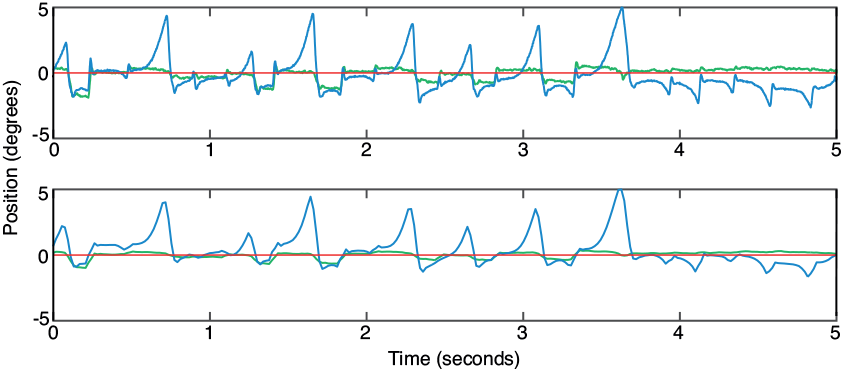
One example of a nystagmus waveform eye movement recording that was subsequently down-sampled and adjusted for the motion-fixation and motion-following conditions. (A) original horizontal (X) position (blue line) and vertical (Y) position (green line) as a function of time. The red line shows the screen centre. (B) X and Y positions for the same waveform after it was down-sampled, adjusted and smoothed. Plotting conventions as in panel A.

### Results

For each participant, stimulus condition (unflanked, horizontal flankers and vertical flankers) and motion condition (no-motion, motion-fixation and motion-following), three blocks of 60 trials were combined to give 180 trials per condition. Within each condition, the proportion of correct responses was determined for each stimulus size presented. Psychometric functions were fitted to the behavioural data for each stimulus condition and motion condition using a cumulative Gaussian function, as in Experiment 1, with thresholds again taken at 62.5% correct.

Figure 6A shows the average gap-size thresholds for all three stimulus conditions (unflanked, horizontal and vertical flankers) in each motion condition, again in both min. of arc and their logMAR equivalents. In the no-motion condition, thresholds were low for the unflanked target (green bar) with slight elevations in thresholds with crowding when the horizontal and vertical flankers were introduced, similar to the control group thresholds in Experiment 1. In the motion-fixation condition, unflanked thresholds were elevated relative to the no-motion condition, though not to the level of the nystagmats in Experiment 1 (dashed lines). In contrast, thresholds for the crowded stimulus conditions were elevated to a level greater than the nystagmus group thresholds, though lower than the amblyopes. In the motion-following condition (where the participant was allowed to follow the stimulus as it moved) we find again a small elevation in unflanked thresholds, with further elevation in the flanked conditions. These crowded elevations were of a lower magnitude than those of the motion-fixation condition, though comparable to the level of performance seen for nystagmats in Experiment 1. Importantly, we see a similar horizontal-elongation of crowding thresholds for both motion-fixation and -following conditions, as for the nystagmats in Experiment 1.

**Figure 6.**
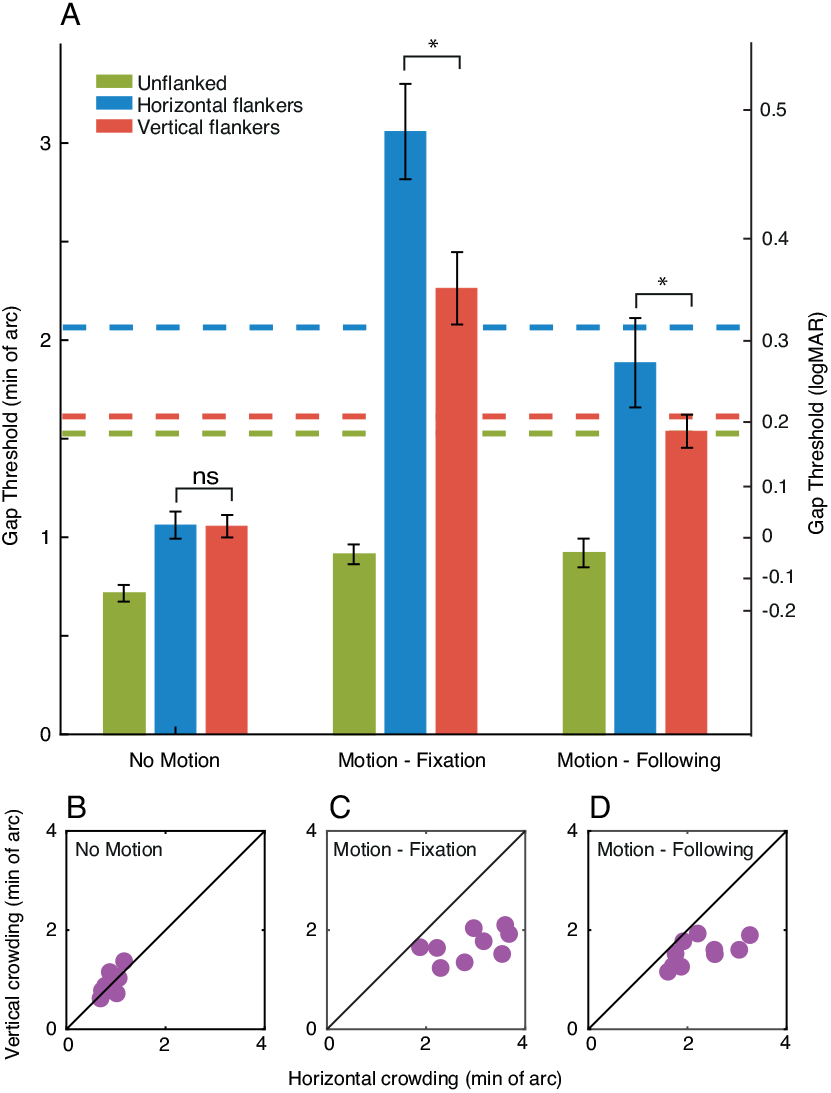
A. Gap thresholds for Experiment 2, with stimulus conditions (unflanked, horizontal and vertical flankers) tested in three motion conditions: no motion, motion-fixation and motion-following. The green bars represent the unflanked condition, blue bars represent horizontal flankers and the red bars represent vertical flankers. Gap thresholds are shown on the left-hand y-axis in minutes of arc and on the right-hand axis with corresponding logMAR values. Dashed coloured lines plot average thresholds for the nystagmus participants for each condition in Experiment 1. *= significant and ns = not significant, error bars represent the SEM. B-D. Horizontal flanker gap thresholds for each individual plotted on the x-axis against vertical flanker gap thresholds on the y-axis, separately for the 3 motion conditions. The black line represents perfect correspondence between horizontal and vertical flanker thresholds. Purple circles represent individual participants, note the individual X and Y scales.

A 3×3 repeated measures ANOVA was performed on thresholds with factors for stimulus condition and motion condition. We find significant main effects of stimulus condition F_(2,18)_ = 109.19, p<0.001, and motion condition F_(2,18)_ = 102.06, p<0.001, and a significant interaction F_(4,36)_ = 22.56, p<0.001. To determine the effect of motion on the horizontal-elongation of crowding we undertook paired sample t-tests between horizontal and vertical flanker conditions. As in Experiment 1, the difference between the horizontal and vertical flanker conditions in the no-motion condition was non-significant, t_(9)_ = 0.123, p=0.905. This difference was however significant in the motion-fixation condition, with horizontally placed flankers producing more crowding than vertically placed flankers, t_(9)_ = 6.481, p<0.001, and likewise in the motion-following condition t_(9)_ = 6.148, p<0.001. This is seen in Figure 8B-D, where the black diagonal line represents the line of unity between thresholds with horizontal and vertical flankers. In the no-motion condition all subjects cluster around the unity line. In contrast, for the motion-fixation and -following conditions we see a clear shift towards higher thresholds with horizontal flankers as all participants lie below the unity line.

To directly compare performance between the nystagmats in Experiment 1 and the simulated conditions in Experiment 2, we next performed a set of independent t-tests. When compared with the motion-fixation control condition, nystagmats had significantly worse thresholds for the unflanked condition t_(16)_= 3.004, p=0.008, whereas in the horizontal flanked condition the nystagmats were significantly better t_(16)_= −2.965, p=0.009. Thresholds were not significantly different with vertical flankers t_(16)_= −1.043, p=0.313. When compared to the motion-following condition, nystagmats again had significantly higher thresholds for the unflanked condition t_(16)_= 3.187, p=0.006, though neither the horizontal t_(16)_= −0.765, p=0.456 nor the vertical t_(16)_= 0.109, p=0.915 flanker conditions differed significantly between the groups. This shows that the motion-following condition in controls produced similarly elevated thresholds to those found with nystagmats when flankers were present, though acuity in the controls remained better than the nystagmats in all cases.

Altogether, the introduction of nystagmic waveform motion to the stimulus impaired performance in adults with typical vision. We also find larger elevations in crowding thresholds with horizontal flankers compared to vertically placed flankers when nystagmus motion is applied to the stimulus, similar to that found in the nystagmus group in Experiment 1. This horizontal-elongation of crowding was significant in both the motion-fixation and motion-following conditions, though overall performance levels were better matched between nystagmats and control participants in the motion-following condition. These findings of a horizontal-elongation of crowding support the hypothesis that the image motion derived from eye movements could be the cause of nystagmic crowding. To probe this further, we next compared properties of the image motion and associated eye movements (stimulus position, velocity, and periods of foveation) with the elevations in crowding thresholds.

#### Eye movement analyses

We first consider the broad properties of the eye movements made by participants in this experiment. Consistent with task requirements, the average variability of horizontal eye positions was greatest in the motion-following condition (0.12°±0.02°), where subjects followed the stimulus, compared to the no-motion (0.08°±0.01°) and motion-fixation (0.09°±0.04°) conditions where fixation had to be maintained. Mean horizontal eye velocity was also greatest in the motion-following condition (7.69°/sec ±0.67°/sec) and lower in the no-motion (6.30°/sec ±0.37°/sec) and motion-fixation (5.94°/sec ±0.71°/sec) conditions. Variability in vertical eye position was similar across the motion conditions: no motion (0.12°±0.01°), motion-fixation (0.11°±0.01°) and motion-following (0.14°±0.02°). Mean vertical eye velocity was lowest in the motion-fixation condition (6.63°/sec ±0.28°/sec), whereas the no-motion and motion-fixation condition were similar at 7.37°/sec ±0.41°/sec and 7.73°/sec ±0.42°/sec respectively. In other words, the motion-following condition generated a bias towards eye movements in the horizontal plane, which was not present in the motion-fixation and no-motion conditions.

To fully account for the differences between these conditions, we also need to incorporate the stimulus motion into calculations. We first determined the absolute difference between the eye position and the stimulus throughout each trial. This was calculated by subtracting the stimulus position from the recorded eye position in both the horizontal and vertical plane. As expected, we find a low positional offset between the eye and stimulus in the horizontal plane for the no-motion condition (0.35°±0.02°), which increased once motion was applied in the motion-fixation (1.36°±0.01°) and motion-following (1.42°±0.01°) conditions. The higher positional offset in the motion conditions is consistent with the elevations in thresholds relative to the no-motion condition, though this analysis cannot explain the differences in performance between the motion conditions.

We next sought a more fine-grained analysis of the relationship between the stimulus and fixation using analyses of foveation duration, as in Experiment 1. In the course of these analyses, we noted a variability in the duration of trials across the three motion conditions. Average trial duration was longest in the motion-following condition (1.75±0.03sec), which decreased in the motion-fixation condition (1.37±0.01sec) and further again in the no-motion condition (1.05±0.01sec). The longer duration of trials in the motion-following condition could partly account for the observed improvements in both crowding tasks compared to the motion-fixation condition.

In order to examine the duration of these trials where fixation fell within the foveation window, we again applied the spatial and velocity parameters from the NAFX to determine the foveation windows for position (±0.5-4°) and velocity (4-10°/sec). Figure 7A shows the average duration of foveation for a range of spatial windows from the NAFX, with the velocity parameter fixed at 10°/sec. As the spatial window widens from 0.5° to 4°, the duration of foveation increases in all three motion conditions. At the smallest spatial windows, foveation periods were longer in the no-motion condition compared to the two motion conditions. By increasing the spatial window of foveation, the motion conditions diverge, with motion-following having longer durations of foveation than motion-fixation. The longer periods of foveation for the motion-following condition with these intermediate spatial windows suggests that participants were indeed able to gain closer foveal proximity to the stimulus in periods of slower eye movements than in the motion-fixation condition where eye movements were not allowed, though neither motion condition reached the degree of foveation possible in the no motion condition (seen in particular at the narrowest windows). As in Experiment 1, we find little change in the average duration of foveation as velocity windows were widened (Figure 7B, where the spatial parameter was set at 4°).

**Figure 7.**
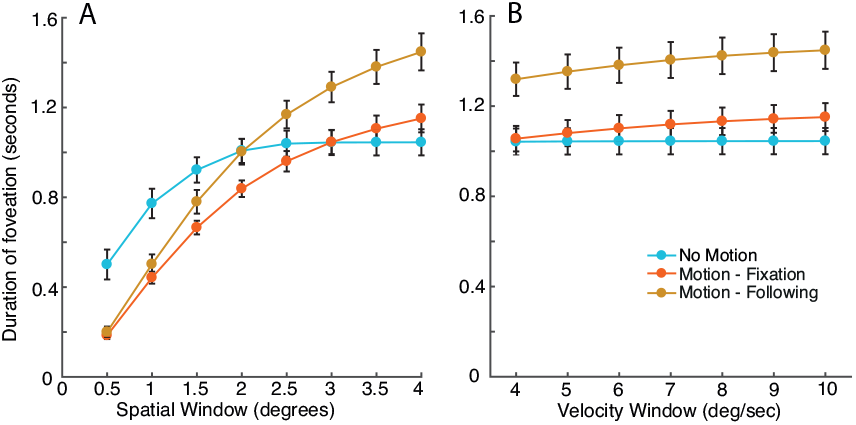
A. Average amount of foveation time across trials where the position of the eye falls within a spatial window around the target for each of the motion conditions (with a fixed velocity window). The X-axis shows the different spatial windows in 0.5° increments. The Y-axis is the total duration of time within the spatial window. B. The average duration of time spent within the foveation velocity window (with a fixed spatial window). The X-axis shows the different velocity windows in 1°/sec increments and the Y-axis is the total duration of the trial foveating. Error bars represent the SEM.

We next divided these trials into best-foveation and worst-foveation as in Experiment 1, described in Appendix D. However, as in Experiment 1, we do not find an effect of foveation on performance in the 3 stimulus conditions. Although we find no differences in performance when splitting the trials for individuals based on foveation, the larger differences in foveation between the motion conditions in Figure 7 are nonetheless consistent with a role for image motion in crowding.

Overall, the results of Experiment 2 demonstrate that the application of nystagmus waveform motion to Landolt-C stimuli can produce a pattern of thresholds in adults with typical vision that is similar to that found in nystagmic crowding, both in terms of overall threshold elevation and the horizontal-elongation of crowding. This is likely due to a reduction in foveation duration, which would both cause both image smear and/or the relocation of the stimulus to more parafoveal locations.

## Discussion

Our aim was to investigate the elevation in visual crowding associated with idiopathic infantile nystagmus, and whether it is the product of a long-term sensory deficit or due to the instantaneous image motion caused by the involuntary eye movements. In Experiment 1, we demonstrate foveal elevations in Landolt-C acuity and crowding for the nystagmus group compared to typical adults, consistent with previous findings (Chung & Bedell, 1995; Pascal & Abadi, 1995). In the nystagmus group we further observe higher thresholds when flankers were positioned horizontally than vertically, an effect that was absent for both typical adults and those with strabismic amblyopia. This horizontal-elongation of crowding in nystagmus is consistent with an origin in image motion caused by the pattern of eye movements. The lack of this effect in amblyopia replicates the finding of Levi and Carney (2011), suggesting a distinct origin for amblyopic crowding that is consistent with broader evidence for an amblyopic sensory deficit (Kiorpes & McKee, 1999). In Experiment 2, we reproduced the horizontal-elongation of crowding in adults with typical vision using stimulus motion derived from the eye movements of participants with nystagmus. This is again consistent with an origin for nystagmic crowding in image motion, unlike crowding effects in typical and amblyopic vision. We propose that the incessant eye movements of nystagmus cause either momentary image smear or the relocation of the stimulus eccentrically into peripheral retina.

Our conclusion that nystagmic crowding results from instantaneous eye motion is consistent with the results of Pascal and Abadi (1995), who found that crowding elevations were clearer in cases of idiopathic nystagmus than in those derived from albinism (despite the latter having higher mean elevations). They attributed the idiopathic crowding elevations to the greater fixational instability and shorter periods of foveation in the idiopathic group. Accordingly, the idiopathic nystagmats tested here in Experiment 1 showed clear elevations in crowded thresholds, with a predominance of jerk-type nystagmus that gave only brief durations of foveation relative to those in the amblyopia and control groups. Although individual differences in foveation and other eye-movement parameters did not correlate with crowded performance (see Appendices B-D), these differences were small relative to those between groups in Experiment 1 and between conditions in Experiment 2. It may be that eye movements must reach a threshold level of variability, magnitude, or velocity before elevations in crowding arise. Group-level differences in the duration of foveation also followed the different levels of crowding observed in typical adults with simulated nystagmus when fixating vs. following the stimulus (Experiment 2). This is consistent with the nystagmus simulations of Chung, LaFrance, and Bedell (2011), which demonstrate the importance of foveation periods in achieving good acuity with unflanked elements.

Consistent with our findings, Chung and Bedell (1995) also found elevations in crowding for control participants with simulated nystagmus. However, these elevations did not reach the same level as participants with nystagmus, which was taken as evidence for a long-term sensory deficit. In contrast, in Experiment 2 of the present study we replicated both the magnitude and pattern of nystagmic crowding (i.e. the horizontal-elongation of crowding) found in participants with nystagmus. It is possible that the type of motion applied to the stimuli could account for this difference. Chung and Bedell (1995) used idealised, repetitive waveforms that may have allowed participants to anticipate the stimulus movement, whereas our use of directly measured nystagmus waveforms gave stimulus motion of greater variability that was clearly more disruptive to the crowded task. This allowed us not only to replicate the magnitude of these elevations in typical adults, but also the horizontal-elongation of crowding, which follows the predominantly horizontal waveform of nystagmic eye movements. Similarly, Bex, Dakin, and Simmers (2003) found little effect of motion on acuity and crowding in the typical peripheral visual field with stimuli that rotated around an iso-eccentric arc in a predictable manner. Falkenberg, Rubin, and Bex (2007) have also demonstrated that measures of acuity and crowding in typical adults were unaffected by high levels of image instability, with random jitter used to mimic the fixation instability seen in patients with low vision. However, these levels of instability were lower than the variability in eye position seen in the nystagmats tested in the present study.

The horizontal-elongation of crowding that we observe in nystagmus participants in Experiment 1 is consistent with performance anisotropies found in other tasks. Greater elevations in contrast detection and grating acuity have been found for vertical than horizontal gratings, in line with the disruptive effect of image smear along the axis of stimulus modulation (Abadi & Sandikcioglu, 1975; Abadi & King-Smith, 1979; Dunn et al., 2014). Bisection acuity thresholds for participants with nystagmus have also been found to be worse for horizontal bisection acuity than vertical at small spatial separations (Ukwade & Bedell, 2012), again consistent with image smear disrupting performance along the axis of stimulus modulation. Our observation of the horizontal-elongation of crowding with simulated nystagmus eye movements in typical vision is also consistent with previously simulated anisotropies. For instance, Ukwade and Bedell (2012) replicated their finding of an anisotropy for bisection acuity with repetitive simulated image motion.

Several studies have however reported performance anisotropies under conditions of brief presentation where image smear would be less disruptive to the task (Abadi & King-Smith, 1979; Ukwade, Bedell, & White, 2002; Dunn et al., 2014). These findings suggest that at least some degree of sensory deficit may arise in nystagmus. Consistent with these impairments for judgements of a single element, our nystagmats showed elevated acuity levels with the unflanked Landolt-C in Experiment 1. However, simulated nystagmus with typical adults in Experiment 2 was insufficient to match this elevation in the unflanked condition, meaning that we cannot exclude the possibility of a sensory deficit that affects the acuity of the nystagmats in our study. In contrast, simulated nystagmus reproduced both the magnitude and anisotropy of nystagmic crowding. Image motion alone is therefore sufficient to explain the crowded deficits in nystagmus. This distinction could arise from a sensory deficit that disproportionately affected early cortical regions – given that crowding is more strongly linked with higher cortex (Anderson et al., 2012), this could lead to elevations in acuity with little-to-no effect on crowding. Alternatively, it is possible that eye movements are more disruptive to multi-element displays than the single-element displays in acuity tasks, given the greater propensity for flankers to smear into the target.

We have argued that nystagmic crowding results from image motion. However, there are several aspects to image motion that may produce this deficit. One aspect of nystagmic image motion that could give rise to crowding is the image smear caused by the eye movements. This smearing of target and flankers into each other as the stimulus moves across the retina would hinder the ability to identify the orientation of the gap in the target Landolt C. More broadly, Nandy and Tjan (2012) have suggested that image smear may be the basis for the radial elongation of crowded interference zones in the typical periphery. However, although individual differences in crowding show some similarity with eye-movement patterns, dissociations are also evident that suggest independence of the two processes in typical vision (Greenwood et al., 2017). Individuals with nystagmus also do not report the perception of image smear, unlike typical adults when nystagmus is simulated (Bedell & Bollenbacher, 1996), suggesting that the role of smear in nystagmic crowding may be limited.

If image smear were to produce crowding-like effects, the interference from the flankers in this case would likely be more closely linked with masking, the impairment of target discriminability by another spatially-overlapping pattern (Campbell & Kulikowski, 1966; Breitmeyer & Ganz, 1976). Crowding and masking show distinct properties – for instance, the critical spacing for masking is proportional to the size of the target whereas crowding is size invariant (Pelli, Palomares, & Majaj, 2004). Masking also impairs both detection and identification of the stimuli, whereas crowding impairs only identification (Pelli, Palomares, & Majaj, 2004). Although crowding can be distinguished from masking in both peripheral vision (Levi, Hariharan, & Klein, 2002b) and amblyopia (Levi, Hariharan, & Klein, 2002c), the same may not be true for nystagmic crowding if image smear were the sole basis for these deficits. Accordingly, Chung, Levi, and Bedell (1996) observed that elevations in a vernier acuity task during motion were largely invariant to stimulus visibility, suggesting that masking due to image smear is unlikely to be a primary factor, at least for deficits related to acuity.

An alternative explanation of the effect of image motion on nystagmic crowding is a change in stimulus location. Given that nystagmus eye movements are predominantly horizontal (Abadi & Bjerre, 2002), their effect would be to shift the stimulus to more peripheral locations along the horizontal meridian where elevations in crowding are larger than the fovea (Bouma, 1970). The radial/tangential anisotropy of peripheral vision would also produce stronger crowding in the horizontal dimension for these locations (Toet & Levi, 1992). For our simulation of nystagmus in Experiment 2, thresholds were higher and more anisotropic in the motion conditions compared to the no-motion conditions. Our analysis of foveation showed that both motion conditions reduced the duration of foveation relative to the no-motion condition, particularly when foveation was defined as a small spatial window. Foveation durations increased as the spatial window was widened, with motion conditions surpassing the no-motion condition for the broadest windows. This demonstrates that the motion conditions caused the stimulus to be viewed in peripheral retina for longer durations. In other words, better performance was obtained in conditions where participants could get closer to the stimulus for longer periods of time. This may also be true for the nystagmats in Experiment 1, who had lower foveation durations than those with typical vision. It is possible that those with nystagmus might base their judgements on the target stimulus when it is in peripheral vision, elevating crowding thresholds and giving rise to the horizontal-elongation of crowding simply due to this peripheral location.

## Conclusions

Overall our study has demonstrated elevated visual crowding in the fovea of participants with idiopathic nystagmus. These crowded elevations were higher with horizontal flankers than vertical flankers (horizontal-elongation of crowding), matching the horizontal bias of the nystagmus eye movements. Both these elevations in visual crowding and the horizontal-elongation of crowding were reproduced in participants with typical vision when stimuli were moved to simulate the pattern of nystagmus. These results, and the dependence of thresholds on foveation duration, suggest that nystagmic crowding is driven by the eye movements relocating the stimulus into peripheral vision, where crowding is known to occur, rather than from a long-term sensory deficit (though we cannot exclude the possibility of a sensory deficit that disrupts acuity). As a consequence, stabilisation of the eye movements in some people with idiopathic nystagmus may provide a benefit to visual function by reducing these visual crowding effects.

## Acknowledgments

The research was funded by Moorfields Eye Charity (GR000074) and the UK Medical Research Council (MR/K024817/1) and supported by the NIHR Biomedical Research Centre at Moorfields Eye Hospital NHS Foundation Trust. We thank Dr Alexandra Kalpadakis-Smith for her help with development of the experiment code.

## Appendices

## Appendix A Clinical characteristics for participants in Experiment 1

**Table 1.** Clinical characteristics of all participants. Key terms: RVA = right visual acuity, LVA = left visual acuity, BEO = both eyes open, NAD = no apparent deviation, L FA = left fully accommodative esotropia, LXT = left exotropia, RCS = right convergent squint, RXT = right exotropia, LCS = left convergent squint, RLR = right lateral rectus, LMR = left medial rectus, BLR = bilateral lateral rectus, FTR = face turn right, FTL = face turn left, HTL = head tilt left. *relates to prior surgery or botulinum toxin (BTXA) treatment to the noted eye muscle.

|  | Age | RVA | LVA | BEO | Cover test | Eye Movements | Head posture | Stereopsis | Nystagmus type |
| --- | --- | --- | --- | --- | --- | --- | --- | --- | --- |
|  | (years) | (logMAR) |  |  |  |  |  | (seconds of arc) |  |
| Nystagmus | 39 | 0.6 | 0.6 | 0.6 | NAD | - | FTR | 300 | Jerk Right |
|  | 19 | 0.3 | 0.2 | 0.2 | NAD | - | FTL | 85 | Pendular |
|  | 26 | 0.5 | 0.5 | 0.5 | NAD | - | FTR HTL | 300 | Jerk Right |
|  | 34 | -0.1 | -0.1 | -0.1 | NAD | - | - | 170 | Jerk Left |
|  | 38 | 0.2 | 0.2 | 0.2 | NAD | - | - | 110 | Jerk Right |
|  | 28 | 0.4 | 0.4 | 0.3 | NAD | - | - | 300 | Pendular |
|  | 20 | 0.6 | 0.6 | 0.6 | NAD | - | - | 85 | Jerk Left |
|  | 38 | 0 | 0 | 0 | NAD | - | FTL | 85 | Jerk Right |
| Amblyopia | 35 | -0.2 | 0.3 | -0.2 | LCS | - | - | 170 | - |
|  | 49 | 0.1 | 0.8 | 0.1 | LXT | - | - | - | - |
|  | 25 | 0.58 | 0 | 0 | RCS | - | - | - | - |
|  | 43 | 0.2 | -0.2 | -0.2 | RXT | -1 RLR* | - | - | - |
|  | 34 | 0.3 | -0.1 | -0.2 | RXT | -0.5 LMR* | - | - | - |
|  | 36 | -0.1 | 0.5 | -0.1 | LCS | -2 LMR* | - | - | - |
|  | 25 | 0 | 0.8 | 0 | LCS L/R | -1 Left Elevation* | - | - | - |
|  | 36 | 0.3 | -0.1 | -0.1 | RCS | - | - | - | - |
|  | 33 | 1 | 0 | 0 | LCS | -1 BLR* | - | - | - |
|  | 46 | 0 | 0.8 | 0 | LXT | - | - | - | - |
| Control | 36 | 0 | 0 | 0 | NAD | - | - | 55 | - |
|  | 34 | 0 | 0 | 0 | NAD | - | - | 55 | - |
|  | 28 | 0 | 0 | 0 | NAD | - | - | 55 | - |
|  | 27 | -0.1 | -0.1 | -0.1 | NAD | - | - | 85 | - |
|  | 26 | 0 | -0.1 | -0.2 | NAD | - | - | 85 | - |
|  | 39 | -0.2 | -0.2 | -0.2 | NAD | - | - | 85 | - |
|  | 21 | -0.1 | -0.1 | -0.1 | NAD | - | - | 85 | - |
|  | 41 | 0.1 | 0.1 | 0.1 | NAD | - | - | 85 | - |
|  | 29 | 0 | -0.1 | -0.2 | NAD | - | - | 85 | - |
|  | 40 | -0.2 | -0.2 | -0.2 | NAD | - | - | 85 | - |

## Appendix B The role of eye movements in Experiment 1

In Experiment 1 of the main paper we describe the features of the overall eye movements in terms of position and speed. We demonstrate that the nystagmus participants have a larger variability in position and with the fastest velocities. If the observed elevations in nystagmic crowding are due to the image motion caused by these eye movements, then properties like variability, velocity, and the duration of foveation should correlate with gap-size thresholds. In other words, with increases in eye position variability and eye speed we should see an increase in the size of gap thresholds.

Figure 8A plots individual gap thresholds against the standard deviation of the horizontal position of the eye for each participant group. The correlation between gap thresholds and the standard deviation of the horizontal eye position was not significant in any testing condition in the control participant group (unflanked r_(9)_=0.042, p=0.921, horizontal flankers r_(9)_=0.22, p=0.596, and vertical flankers r_(9)_=-0.166, p=0.693). This was the same in the amblyopic group (unflanked r_(9)_=-0.134, p=0.711, horizontal flankers r_(9)_=-0.007, p=0.984 and vertical flankers r_(9)_=0.108, p=0.782) as well as the nystagmus group (unflanked r_(7)_=-0.037, p=0.919, horizontal flankers r_(7)_=0.019, p=0.957 and vertical flankers r_(7)_=0.153, p=0.672). Similarly, there was no significant correlation between horizontal eye velocity and thresholds (Figure 8B) in the control group (unflanked r_(9)_=0.144, p=0.734 horizontal flankers r_(9)_=0.282, p=0.498 and vertical flankers r_(9)_=-0.079, p=0.851). This was the same in the amblyopic group (unflanked r_(9)_=-0.082, p=0.820, horizontal flankers r_(9)_=0.397, p=0.256 and vertical flankers r_(9)_=0.047, p=0.895) as well as the nystagmus group (unflanked r_(7)_=0.206, p=0.568, horizontal flankers r_(7)_=0.156, p=0.668 and vertical flankers r_(7)_=-0.196, p=0.588). Finally, there were no significant correlations with gap thresholds for either the variability in vertical eye position or velocity (data not shown). Altogether, there is no association between gap thresholds and either mean velocity or variability in eye position.

**Figure 8.**
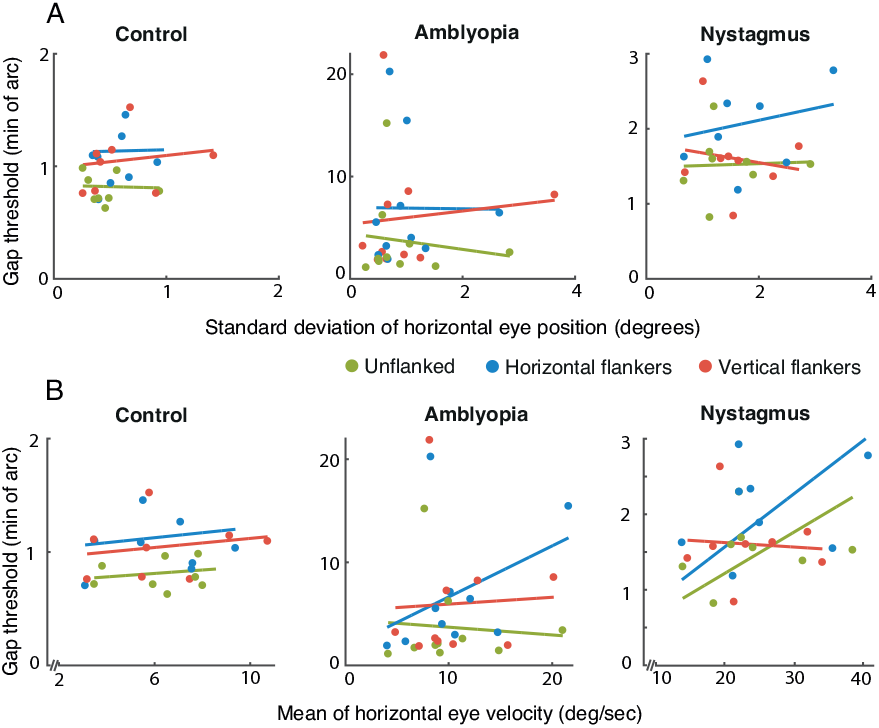
A. Gap thresholds plotted against the standard deviation of the horizontal eye position for each of the stimulus conditions and for each individual in the three participant groups (separate panels). B. Gap thresholds plotted against the mean of the eye velocity in the horizontal plane. Coloured circles separate the data into the three stimulus conditions (unflanked, horizontal and vertical flankers). Coloured lines represent the best fit linear function of the corresponding-coloured data. Note the individual x and y axis scales for each participant group.

## Appendix C The role of foveation in Experiment 1

Here we examine the role of foveation on performance in Experiment 1 by dividing trials into best-foveation and worst-foveation categories and re-fitting psychometric functions to these split datasets. Trials were divided by first examining the distribution of foveation durations to find a combination of spatial and velocity windows that produced a normal distribution of durations across all trials, without any ceiling or floor effects. We found that parameters of ±2° for position and 8°/sec for velocity gave normal distributions for all nystagmus participants. We then applied these foveation criteria to data from all the nystagmic individuals to calculate the foveation duration within each trial. A median split was then applied to an individual’s data to identify trials with best-foveation and worst-foveation, resulting in 120 trials for each. The proportion of correct responses was then calculated for each gap size, separately for the best and worst foveation trials, and new psychometric functions were fit to calculate gap-size thresholds. If nystagmic crowding is caused by image motion, and in particular by foveation, we should find lower gap thresholds for all stimulus conditions (unflanked, horizontal and vertical flankers) and in particular a decrease in the difference between the horizontal flanker and vertical flanker stimulus conditions for best-foveation trials compared to the worst-foveation trials.

Figure 9 shows thresholds obtained from the best-foveation and worst-foveation trials for each stimulus condition in the nystagmus group. As in the main analysis, for each foveation condition, thresholds for the unflanked stimulus were lower than both horizontal and vertical flanker stimulus conditions, with an elevation of the horizontal flanker gap thresholds relative to the vertical flanker gap threshold. We compared our three stimulus conditions (unflanked, horizontal and vertical flankers) during trials with best-foveation and worst-foveation and undertook a 2×3 repeated measures ANOVA, with factors for foveation and stimulus condition. LogMAR values were used as before to reduce heteroscedasticity in the data. The ANOVA showed a significant main effect of stimulus condition F_(2,14)_= 24.634, p<0.001, though the main effect for foveation was not significant, F_(1,7)_=1.949, p=0.205, showing that thresholds were indistinguishable when derived from trials with either best- or worst-foveation. The interaction between stimulus condition and foveation was not significant, F_(2,14)_=0.092, p=0.912, again demonstrating that best-foveation or worst-foveation did not influence the pattern of thresholds between the stimulus conditions. The horizontal-elongation was present in both best- and worst-foveation conditions, with a significant difference between horizontal and vertical flanker conditions in each case (best foveation: t_(7)_=4.698, p=0.002; worst-foveation: t_(7)_=4.005, p=0.005).

**Figure 9.**
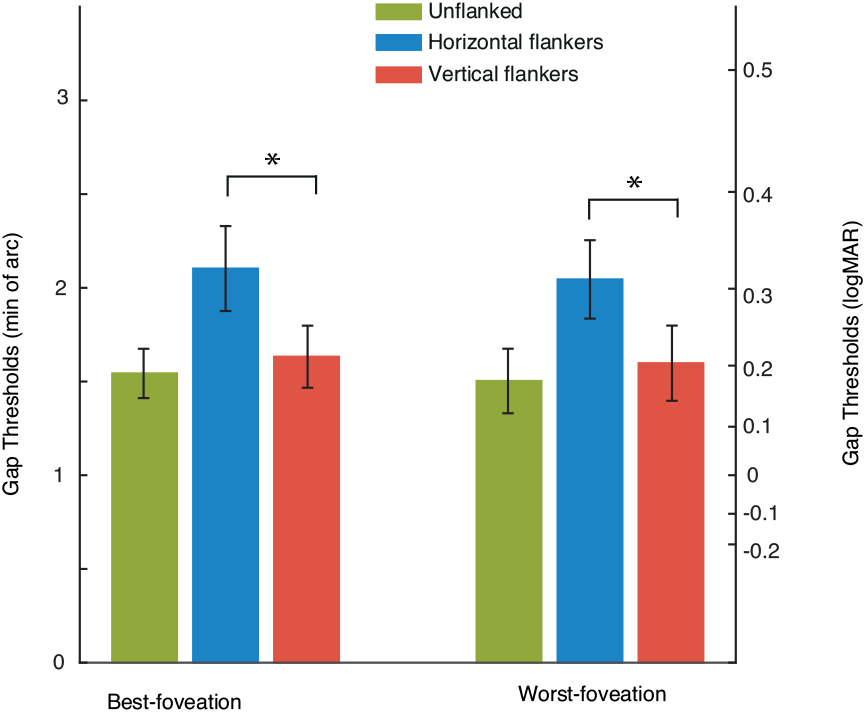
Bar plot showing the average gap thresholds for the nystagmus participants split by best-foveation and worst-foveation trials. The data show similar thresholds for within best-foveation and worst foveation for all stimulus conditions. We plot a secondary y-axis to represent the logMAR equivalent, calculated by taking the logarithm of the mean values. A significant horizontal-elongation of crowding is seen in both best- and worst-foveation conditions. Green bars represent the unflanked stimulus condition, blue bars the horizontal flanked condition and red bars the vertical flanked condition, *= significant and ns = not significant, error bars represent the SEM.

There are two caveats with this analysis, however. First, by splitting the data in half, we reduce the reliability of the psychometric function fits and the gap thresholds derived from these fits. Accordingly, we find a reduction in the goodness-of-fit for the median-split analyses – the squared error between psychometric functions and data (weighted by the number of trials in each point) increases when trials are split into best-(0.056) and worst-foveation (0.043) compared to the full data set (0.030). Thus, although the median split would separate trials according to the degree of foveation, subtle differences in gap thresholds between these foveation conditions may be difficult to identify due to the reduction in data. Second, although these differences in foveation duration cannot account for the horizontal elongation of crowding on an individual basis, the differences in foveation introduced with these median-split analyses are small relative to the larger differences at the group level (shown in Figure 4). It may be that these larger differences in foveation are required to drive the difference in performance observed between groups.

## Appendix D The role of foveation in Experiment 2

We also considered the role of foveation in Experiment 2, using the same methodology as described above in Appendix C. We used the same parameters for position (±2°) and velocity (8°/sec) as the stimulus motion was generated from the nystagmus waveforms measured in Experiment 1. If the elevations in crowding observed in our typical observers in Experiment 2 is caused by image motion generated from the nystagmus waveform motion, then we should find lower gap thresholds for all stimulus conditions (unflanked, horizontal and vertical flankers) in the best-foveation trials.

Figure 10 shows thresholds obtained from the best-foveation and worst-foveation trials for each stimulus and motion condition in Experiment 2. We find that for each foveation condition unflanked thresholds are very similar and lower than both flanked conditions. Thresholds for horizontal- and vertical-flanked conditions were similar in the no-motion condition for both best- and worst-foveations. Similarly, in both motion conditions (motion fixation and motion following) we find higher gap thresholds with horizontal flankers than with vertical flankers, with a similar magnitude regardless of foveation.

**Figure 10.**
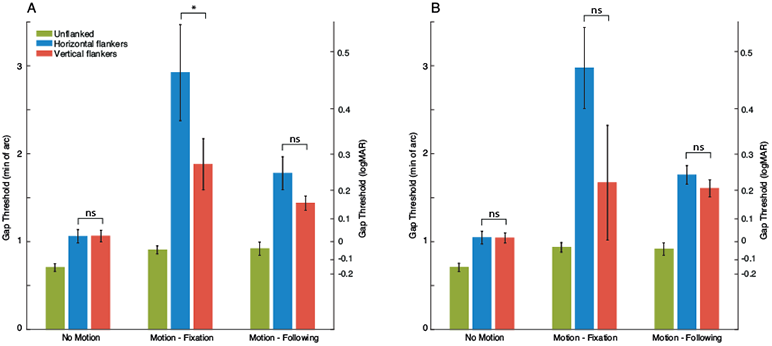
Average gap thresholds for the control participants split by best-foveation (A) and worst-foveation trials (B). Values are plotted in minutes of arc (left y-axis), along with logMAR equivalents (right y-axis). Green bars represent the unflanked stimulus condition, blue bars the horizontal flanked condition and red bars the vertical flanked condition, *= significant and ns = not significant, error bars represent the SEM. The data show similar thresholds for best-foveation and worst foveation in all stimulus conditions, with a significant horizontal-elongation of crowding in both cases.

We compared our three stimulus conditions (unflanked, horizontal and vertical flankers) during trials with best-foveation and worst-foveation and undertook a 2×3×3 repeated measures ANOVA with factors for foveation, motion condition and stimulus condition. LogMAR values were used here to reduce heteroscedasticity in the data. The ANOVA showed a significant main effect of stimulus condition F_(2,18)_= 51.185, p<0.001, indicating that flankers and their location affected thresholds. We found a significant effect of motion F_(2,18)_= 28.788, p<0.001, showing that the motion applied to the stimulus affected thresholds. The main effect for foveation was not significant, F_(1,9)_=0.473, p=0.509, showing that thresholds were not affected by best- or worst-foveation during application of nystagmus motion to the stimulus.

The interaction between motion condition and foveation was not significant, F_(2,18)_=2.498, p=0.110, demonstrating that best-foveation or worst-foveation did not influence the pattern of thresholds between the motion conditions. The interaction between stimulus condition and foveation was also non-significant F_(2,18)_=2.347, p=0.124 indicating that foveation did not alter the pattern of thresholds across stimulus conditions. As in the main analysis, we did find a significant interaction between the motion condition and stimulus condition F_(4,36)_=6.773, p<0.001, signifying the effect of motion on thresholds in the different stimulus conditions. The three-way interaction was however non-significant, F_(4,36)_=2.464, p=0.062, demonstrating again that foveation does not alter the relationship between stimulus and motion conditions.

Altogether, the horizontal-elongation was present in both best- and worst-foveation conditions and with both motion-fixation and -following conditions. However, the difference between thresholds with horizontal vs. vertical flankers was only significant in the motion-fixation condition whilst in best-foveation t_(9)_=2.415, p=0.039. All other comparisons between horizontal- and vertical-flanked conditions were non-significant. As above however, the overall similarity between best- and worst-foveation conditions is likely due to the reduction of data in each condition of the median-split analysis and the resulting increase in error. Additionally, the small differences in foveation duration between median-split conditions may be insufficient to alter the horizontal elongation of crowding, relative to the larger differences induced between fixation conditions (Figure 7).

